# Post-transcriptional regulation of redox homeostasis by the small RNA SHOxi in haloarchaea

**DOI:** 10.1101/2020.10.12.336255

**Authors:** Diego Rivera Gelsinger, Rahul Reddy, Kathleen Whittington, Sara Debic, Jocelyne DiRuggiero

## Abstract

Haloarchaea are highly resistant to oxidative stress, however, a comprehensive understanding of the processes regulating this remarkable response is lacking. Oxidative stress-responsive small non-coding RNAs (sRNAs) have been reported in the model archaeon, *Haloferax volcanii*, but targets and mechanisms have not been elucidated. Using a combination of high throughput and reverse molecular genetic approaches, we elucidated the functional role of the most up-regulated intergenic sRNA during oxidative stress in *H. volcanii*, named **<u>S</u>**mall RNA in **<u>H</u>**aloferax **<u>Oxi</u>**dative Stress (**SHOxi**). SHOxi was predicted to form a stable secondary structure with a conserved stem-loop region as the potential binding site for trans-targets. NAD-dependent malic enzyme mRNA, identified as a putative target of SHOxi, interacted directly with a putative “seed” region within the predicted stem loop of SHOxi. Malic enzyme is an enzyme of the tricarboxylic acid cycle that catalyzes the oxidative decarboxylation of malate into pyruvate using NAD^+^ as a cofactor. The destabilization of malic enzyme mRNA, and the decrease in the NAD^+^/NADH ratio, resulting from the direct RNA-RNA interaction between SHOxi and its trans-target was essential for the survival of *H. volcanii* to oxidative stress. These findings indicate that SHOxi likely regulates redox homeostasis during oxidative stress by the post-transcriptional destabilization of malic enzyme mRNA. SHOxi-mediated regulation provides evidence that the fine-tuning of metabolic cofactors could be a core strategy to mitigate damage from oxidative stress and confer resistance. This study is the first to establish the regulatory effects of sRNAs on mRNAs during the oxidative stress response in Archaea.

## INTRODUCTION

Small non-coding RNAs (sRNAs) are important regulators for multiple cellular functions across the 3 domains of life (1). sRNAs are ubiquitous in Bacteria and Eukarya, playing essential roles in transcriptional regulation, RNA processing and modification, mRNA stability, and translation regulation (1–3). Regulatory sRNAs have also been found in Archaea, but few have been functionally characterized and many questions remain (4–8).

RNA sequencing (RNA-seq) applied to a small number of archaeal species, including *Haloferax volcanii, Haloferax mediterranii, Methanosarcina mazei,* and *Sulfolobus solfataricus*, revealed that hundreds to thousands of sRNAs were potentially transcribed from those gene-dense genomes (9–23). Archaeal sRNAs range from 50 to 500 nucleotides in size and can be categorized into three classes: intergenic sRNAs, antisense sRNAs, and sense sRNAs (9, 14, 17). Molecular studies to uncover the biological roles of these non-coding transcripts have focused on intergenic sRNAs (13, 20, 22). These studies found that archaeal sRNAs can regulate target mRNAs through base pairing and consequently alter stabilization of the target transcripts or mask the ribosome binding site, decreasing translation (13, 20, 22).

Archaeal sRNAs have been implicated in several biological functions such as cellular growth, osmolarity, carbon and energy metabolism, nutrient uptake, stress response, and biofilm formation, which underscores their importance for cellular functionality (9, 11, 13, 22–24). For example, in the methanogen *M. mazei*, sRNA_154_ was up-regulated under nitrogen starvation conditions, affecting multiple targets such as nitrogenase and glutamine synthetase, and sRNA_162_ was shown to regulate the switch between carbon and energy sources (13, 25). Diverse sRNA regulatory mechanisms have also been elucidated in *M. mazei*; sRNA_162_ and sRNA_41_ were reported to bind in *trans* to the ribosome binding site (RBS) of bicistronic mRNAs and to bind in *cis* to the 5’ leader region of another mRNA, decreasing the translation of its targets, while sRNA_154_ was shown to bind multiple targets, affecting the stability of those transcripts (13, 21, 26). A large number of sRNAs have been reported in the halophilic model archaeon, *H. volcanii,* and a few of these have been assigned potential function, including adaptation to phosphate starvation conditions (22), and oxidative stress response (9). However, despite the genetic tools available for *H. volcanii* (27) limited sRNA-dependent regulatory mechanisms have been elucidated.

Oxidative stress occurs when the level of reactive oxygen species (ROS) produced in cells by aerobic metabolic activity or environmental challenges overwhelms antioxidant defense mechanisms and damage accumulates (28). Oxidative stress is universal in all domains of life and often produces robust phenotypes in cells (29). This stressor has profound implications for cell survival, and in humans, oxidative stress plays an important role in aging and many disease states (29, 30). Regulation of the oxidative stress response in halophilic archaea is of particular interest because these organisms are extremely resistant to oxidative stress (9, 31–35). In the haloarchaeon *Halobacterium salinarum,* the transcription factor RosR was found to be highly responsive to changes in oxygen levels and to control the expression of over 300 genes in response to ROS damage (31). This work demonstrated that the oxidative stress response of haloarchaea is highly regulated and mediated, in part, by transcription factors. RosR showed no differential expression during oxidative stress in *H. volcanii*, suggesting that it plays a different role in this organism and that other factors, such as a sRNAs, could be key players in regulating its response to oxidative stress (9). In bacteria, sRNAs have been assigned as key players in the oxidative stress response, protecting cells from ROS in various ways (36). For example, the sRNA OxyS confers genomic stability in *Escherichia coli*, iron metabolism is tuned by the sRNA RhyB in *Salmonella typhimurium*, various transporters are regulated by the sRNAs SorX and SorY to alter metabolism and homeostasis in *Rhodobacter sphaeroides*, and the ROS detoxifying enzyme catalase is controlled by the sRNA OsiA in *Deincoccus radiodurans,* all in response to oxidative stress (36–42). Detailed knowledge of the functions of sRNAs in archaea is limited to only a few examples, and none of these have been implicated in the response to oxidative stress. In a previous sRNA-seq screen, we identified hundreds of sRNAs differentially expressed in response to oxidative stress, including both intergenic and antisense sRNAs (9), providing the opportunity to address the mechanistic and functional role of sRNAs in the oxidative stress response of *H. volcanii*.

Here we applied a combination of high throughput and reverse molecular genetic approaches to determine a mechanism of action for the most up-regulated intergenic sRNA during oxidative stress in *H. volcanii.* This sRNA, previously identified as sRNA STRG.277.2 (9), was found to play an integral role in the regulation of redox homeostasis and the survival of *H. volcanii* during oxidative stress. Based on these findings, STRG.277.2 was re-named **<u>S</u>**mall RNA in **<u>H</u>**aloferax **<u>Oxi</u>**dative Stress (**SHOxi**).

## MATERIAL AND METHODS

### Culture growth conditions

*H. volcanii* auxotrophic strain H53 (*Δpyre2, ΔtrpA*) and H98 (*Δpyre2, ΔthyH*) were used for all experiments. Culturing in liquid and solid media was done in rich medium (Hv-YPC) or selection medium (Hv-Cab), at 42°C and with shaking at 220 rpm (Amerix Gyromax 737) (43). Uracil, tryptophan, thymidine, and hypoxanthine were added to a final concentration of 50 μg/mL, each.

### Knockout mutant generation

Deletion mutants of SHOxi (*ΔSHOxi*) were constructed independently in H53 and H98 strain backgrounds using a pop-in pop-out method previously described (44). 500 base pairs upstream and downstream of SHOxi, including small overhangs (30 bp), were PCR amplified, stitched together, and then cloned into the integration vector pTA131 to build the knockout plasmid construct. *H. volcanii* strains H53 and H98 were transformed with the plasmid to yield pop-in clones by uracil autotrophy. To generate the pop-out strains, cells were plated on medium containing 5-fluoro-orotic acid (5-FOA). Deletions were verified at the DNA level by PCR and at the RNA level by northern blot and RNA-seq.

### Oxidative stress exposure

*H. volcanii* liquid cultures were exposed to H_2_O_2_ as previously described (9). In brief, cultures were grown in 160 mL of Hv-YPC or Hv-Cab under optimal conditions to an OD of 0.4 (mid exponential phase). 2 mM H_2_O_2_ was directly added to the cultures followed by an hour incubation at 42 °C with shaking at 220 rpm. Cultures were then rapidly cooled down, centrifuged at 5,000 x g for 5 minutes and the pellets resuspended in 18% sea water. The cell suspensions were then transferred to a 1 mL tube and centrifuged at 6,000 x g for 3 minutes, the pellets were flash frozen and stored at −80 °C until ready for RNA extraction.

### RACE analysis

The 5’ and 3’ ends of SHOxi were determined using the Takara SMARTer Rapid Amplification of cDNA Ends (RACE) kit with slight modifications on total RNA extracts from oxidative stress treated cells. For 5’ RACE the protocol for cDNA generated by random primers was used, followed by the standard protocol with custom internal reverse primer complementary to SHOxi. For 3’ RACE, total RNA was treated with polyA polymerase (NEB) for 1h at 37 °C to add polyA-tails to RNAs. Afterwards, the standard 3’ RACE protocol was followed.

### Overexpression experiments

A variant of the overexpression plasmid pta1228 (45) was built to prevent the introduction of an ATG at the beginning of SHOxi. Using the standard protocol of the Q5 Site-directed Mutagenesis kit (NEB), the region of pTA1228 spanning restriction sites *NdeI* and *BamHI* was replaced with *KasI* yielding the new plasmid pTA1300. The full length of SHOxi was PCR amplified with overhangs and sticky end ligated at the *KasI* restriction site of pTA1300. *H. volcanii* was transformed using the pop-in method and uracil autotrophy to generate *ΔSHOxi* overexpression clones in both H53 and H98 backgrounds. Overexpression was induced at OD 0.4 by addition of 2mM tryptophan and incubation at 42 °C with shaking at 220 rpm for 1 hour. Cells were harvested as described above and used for mRNA-seq or qPCR.

### Oxidative stress survival and growth curves

Assessment of survival in *H. volcanii* wild type and ΔSHOxi under acute oxidative stress conditions (2 mM H_2_O_2_) was done using microdilution plating as described in (9). Counts were averaged and standard deviation calculated between replicates. Survival was calculated as the number of viable cells following H_2_O_2_ treatment divided by the number of viable untreated cells and graphed with standard error bars. Growth curves were done by measuring OD_600_ over time intervals of the wild type and ΔSHOxi exposed to chronic oxidative stress (500 μm H_2_O_2_).

### RNA extraction

Total RNA was extracted using the Zymo Quick-RNA Miniprep kit with the following modifications: *H. volcanii* liquid culture is slimy and viscous thus to increase cellular lysis a 23 G needle and syringe were used to break down the cell pellet after addition of RNA lysis buffer to the frozen pellets to ensure complete cell lysis. Total RNA was then extracted following the standard kit protocol.

### Messenger RNA-sequencing library preparation (mRNA-seq)

Total RNA was DNase I (NEB) treated (37 °C for 2 hours) as previously described (9). Total RNA was then rRNA-depleted using the Ribo-zero Bacteria kit (Illumina). Strand-specific libraries were prepared using the SMART-seq Ultralow RNA input kit (Takara), insert sizes checked with the Bioanalyzer RNA pico kit (Agilent), and either paired-end sequenced (2 × 150 bp) or single-end sequenced (100 bp) on the Illumina HiSeq 2500 platform at the Johns Hopkins University Genetic Resources Core Facility (GRCF).

### mRNA-seq differential expression analysis

We used a read count-based differential expression analysis to identify putative targets of SHOxi that were differentially expressed during oxidative stress and in *ΔSHOxi*. The program featureCounts was used to rapidly count reads that map to the NCBI *H. volcanii* annotation. featureCounts was run with strand-specific options on, paired-end mode on or off, multi-mapping off. The read counts were then used in the R differential expression software package DESeq2 (46). Briefly, read counts were converted into a data matrix and normalized by sequencing depth and geometric mean. Differential expression was calculated by finding the difference in read counts between the SHOxi knockout oxidative stress state to the normalized read counts from the wild-type oxidative stress normalized read counts. The differentially expressed mRNAs were filtered based on the statistical parameter of False Discovery Rate (FDR) under 5%. In addition, only mRNAs with converse differential expression levels (FDR < 5%) in our previous wild type no challenge/oxi stress differential expression comparison (9) were labeled as specific putative targets of SHOxi.

### Northern Blot analysis

20 μg of total RNA and P^32^ ATP end-labeled Century+ RNA markers were loaded onto 5% denaturing urea polyacrylamide gels (SequaGel, National Diagnostics) and run at 30 watts for 1.5 hours to ensure well-spaced gel migration from 50 to 1,000 nucleotides (nt). Gels were transferred onto Ultra-hyb Nylon membranes and hybridized with probes. For SHOxi, we probed with [γ-P^32^] dATP randomly primed amplicons generated with custom primers. Probe primers were at a minimum 10 nt inwards from the predicted genomic coordinates (start and stop) to ensure accurate transcript detection. Hybridizations were done at 65°C. The rpl30 protein (HVO_RS16975) transcript was used as a loading control for differential expression calculation because it was not differentially expressed under oxidative stress in our previous RNA-seq dataset. Differential expression was calculated using ImageJ.

### In silico RNA interactions

The program IntaRNA (47) was used to computationally predict possible interactions with SHOxi and all RNAs in the NCBI *H. volcanii* gene annotation. Options used were no-seed, 42 °C temperature, no START. Top candidates were the top 100 hits ranked by lowest p-value.

### RNA half-life measurement

Wild type or *ΔSHOxi* cells at OD 0.4 were grown for 30 min with H_2_O_2_ to induce endogenous expression of SHOxi and subsequently treated with 100 μg/ml actinomycin D to inhibit transcription. Samples were harvested at 0,15,30, and 60 minutes post-actD, extracted for RNA, and malic enzyme mRNA levels were measured with qRT-PCR at 0, 15, 30, and 60 minutes post-actD, in *ΔSHOxi* and wild type, under oxidative stress.

### Binding site mutagenesis experiments

Mutations within the stem-loop binding site of SHOxi were constructed using the Q5 Site-directed Mutagenesis kit (NEB) standard protocol on the previously described SHOxi overexpression construct (in pTA1300). The forward primer (5’-CCGACACACGGCGTCGCGGTGCGGCCCCCCT-3’) and reverse primer (5’-CGGACTGGCCGACGCCCC-3’) were annealed at 78°C and overhangs were used to introduce point and di-nucleotide mutations through inverse PCR. The mutated SHOxi constructs were verified by Sanger sequencing (Genewiz). The empty vector pTA1300 and the various mutant SHOxi constructs were transformed into both H53 and H98 *ΔSHOxi H. volcanii* strains, as previously described. Overexpression of the mutant SHOxi transcripts were induced under no challenge conditions with 2mM tryptophan for 1h and harvested for RNA extraction and cDNA generation as described in the overexpression experiments. Malic enzyme mRNA expression was then measured via qPCR (SYBR Green PCR Master Mix, ThermoFisher) using primers for malic enzyme and rp130 as a housekeeping gene.

### Dinucleotide luciferase assay

All dinucleotides (NAD^+^, NADH, NADP^+^, NADPH) were extracted using a custom protocol from Promega. In brief, 300μl high pH Bicarbonate Buffer + 1% DTAB was added to cell pellets to lyse the cells. The lysate was split into two 100 uL aliquots, where one was treated with 100μl 0.4N HCl to one tube for acid treatment. Both aliquots were heated at 60°C for 15 min and then cooled at RT for 10 min. The acid treated aliquot was then neutralized with 100μl 0.5M Trizma Base to get oxidized forms of NAD and NADP. The base-only treated sample was neutralized with 200μl 0.4NHCl/0.5M Trizma Base to get reduced forms of NADH and NADPH.

After dinucleotide extraction, extracts were used in the corresponding Glo^TM^ Assay (Promega) using the standard protocol where 50 uL of extract was added with 50 uL of Glo^TM^ Detection Reagent (Promega) in a white bottom 96 well plate (Corning). After 30 minutes of incubation the plates were measured for luminescence using a Glomax Navigator with dual injection pumps, model GM2010 luminometer.

### Protein carbonyl western blotting

Cells were treated and pelleted as previously described. For protein extraction, frozen pellets were resuspended in 1 mL ice cold 1M salt buffer (50 mM potassium phosphate pH 7.0, 1M NaCl, 1% 2-mercaptoethanol) and sonicated 30 seconds ON/ 30 seconds OFF for 30 minutes at room temperature. Lysates were centrifuged at 12,000 x g for 30 minutes at 4°C and the supernatant transferred into a new tube and kept on ice. Protein concentrations were measured using the Quick start Bradford 1x assay standard protocol (Bradford). Protein carbonyls were measured by Western blotting using the OxyBlot kit standard protocol. Briefly, 20 ug of proteins were derivatized with DNPH and ran on a 4-20% SDS PAGE at 120 V for 30 mins. Proteins were transferred to 0.2 uM PVDF membrane (Ambion) in a Trans-blot Turbo (BioRad) for 7 minutes and incubated with primary and secondary antibodies for 1 hour at room temperature, each. ECL+ reagent was added to the blots, and incubated at room temperature, and the blots were scanned with a Typhoon phosphoimager.

### RNA-seq data

All raw reads and processed data from these experiments are available at the National Center for Biotechnology Information under GEO accession number GSE138990.

## RESULTS

### SHOxi is a small non-coding RNA highly responsive to oxidative stress

We previously carried out a sRNA-seq screen in *H. volcanii* under no challenge and oxidative stress conditions and found thousands of differentially expressed sRNAs (9). In this screen, we found a novel transcript, STRG.277.2, with no coding capacity (**Fig. S1**). STRG.277.2 was enriched 21-fold under oxidative stress conditions (2 mM H_2_O_2_ exposure for 1 hour; 80% survival) (**Fig. 1A**), making it the most up-regulated sRNA in our data set (9). We validated with Northern blot analysis that STRG.277.2 sRNA was highly expressed under oxidative stress, with only low levels present under no challenge conditions (**Fig. 1B**). The sequence of this sRNA had a high GC content (87%) compared to the average GC content of the *H. volcanii* genome (61.1%). Its gene was located in an intergenic region of the main chromosome and was flanked by two genes encoding hypothetical proteins on the opposite strand (**Fig. 1C**). We identified the transcription start site (TSS) and transcription termination site (TTS) of STRG.277.2 sRNA with 5’- and 3’-RACE, resulting in a native sRNA of 234 nt in size (Chromosome: NC_013964.1, start: 145098, stop: 145333, strand: minus). The experimental TSS of STRG.277.2 sRNA matched the TSS predicted from our RNA-seq assembly and allowed for accurate characterization of basal archaeal transcription factor binding motifs, including the B recognition element (BRE), the TATA box, the initially melted region (IMR), and the initiator element (Inr) upstream of the TSS (**Fig. 1C**).

**FIGURE 1:**
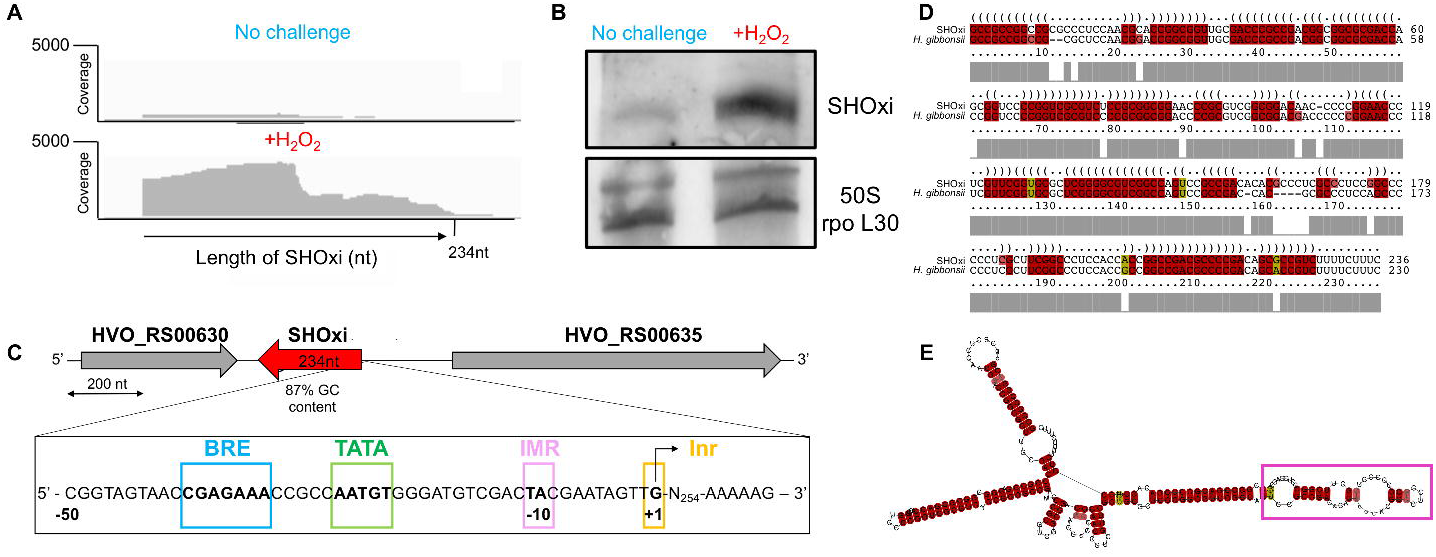
Characterization of SHOxi. (A) RNA-seq coverage plots of the assembled SHOxi transcript with data from (9) under no challenge and oxidative stress conditions. (B) *In vivo* validation of SHOxi expression by Northern blot analysis. 50S rpo L30 was used as a loading control. (C) Genomic context of SHOxi. The inlet box is 50nt upstream of the transcription start site of SHOxi and marked are various conserved archaeal transcription motifs. (D) Multiple sequence and structural alignment of SHOxi and the sRNA homolog in *H. gibbonsii*. (E) Predicted secondary structure model (minimum free energy) of SHOxi. The color indicates number of base pair types (red: 1, yellow: 2), hue shows sequence conservation of the base pair, and saturation indicates the structural conservation of the base pair. Putative interaction region highlighted in magenta.

Using blastn search and the NCBI nt database (version 2019/03/22), we found one highly conserved homolog of SHOxi in *Haloferax gibbonsii,* one of 6 *Haloferax* genomes publicly available. The *H. gibbonsii* sequence was 94% identical at the nucleotide (nt) level, including the upstream regulatory regions, and was located in an intergenic region flanked by two genes with similar predicted functions than those in *H. volcanii*. Because of its drastic response to oxidative stress and its conservation, we named the STRG.277.2 sRNA **<u>S</u>**mall RNA in **<u>H</u>**aloferax **<u>Oxi</u>**dative stress, or SHOxi (referred to as such from here on out).

Using SHOxi and its homolog in *H. gibbonsii*, we generated a multiple sequence alignment with LocARNA and assessed structural conservation (Fig 1D). We predicted a stable secondary structure (**Fig. 1E**) containing high sequence reliability in the first 100 nt and high structural reliability in the last 100 nt, with small drops in reliability in between corresponding to potential loop regions (**Fig. S2**). Although highly structured due to extensive GC base pairing, SHOxi was predicted to form loops and stem loop regions available for base pairing with mRNA targets (**Fig. 1E**).

### SHOxi alters survival during oxidative stress

To assess whether SHOxi played a physiological role during oxidative stress in *H. volcanii,* we knocked out the transcriptional loci, using a pop-in pop-out method previously established (44), and generated a deletion mutant (*ΔSHOxi*). We confirmed the genomic deletion using PCR and the absence of transcript using Northern blot analysis and RT-PCR (**Fig. 3A**, **Fig. S3**). We found no significant difference in the growth rate of the SHOxi deletion mutant when compared to wild type (WT) under no challenge condition or under chronic oxidative stress (500 μm of H_2_O_2_) (**Fig. S4**). However, when *ΔSHOxi* was exposed to 2 mM H_2_O_2_ for 1 hour, replicating the oxidative stress conditions from our sRNA-seq screen (9), we found a severe decrease in survival (avg. 22% survival) when compared to WT (avg. 78% survival) (**Fig. 2A**).

**FIGURE 2:**
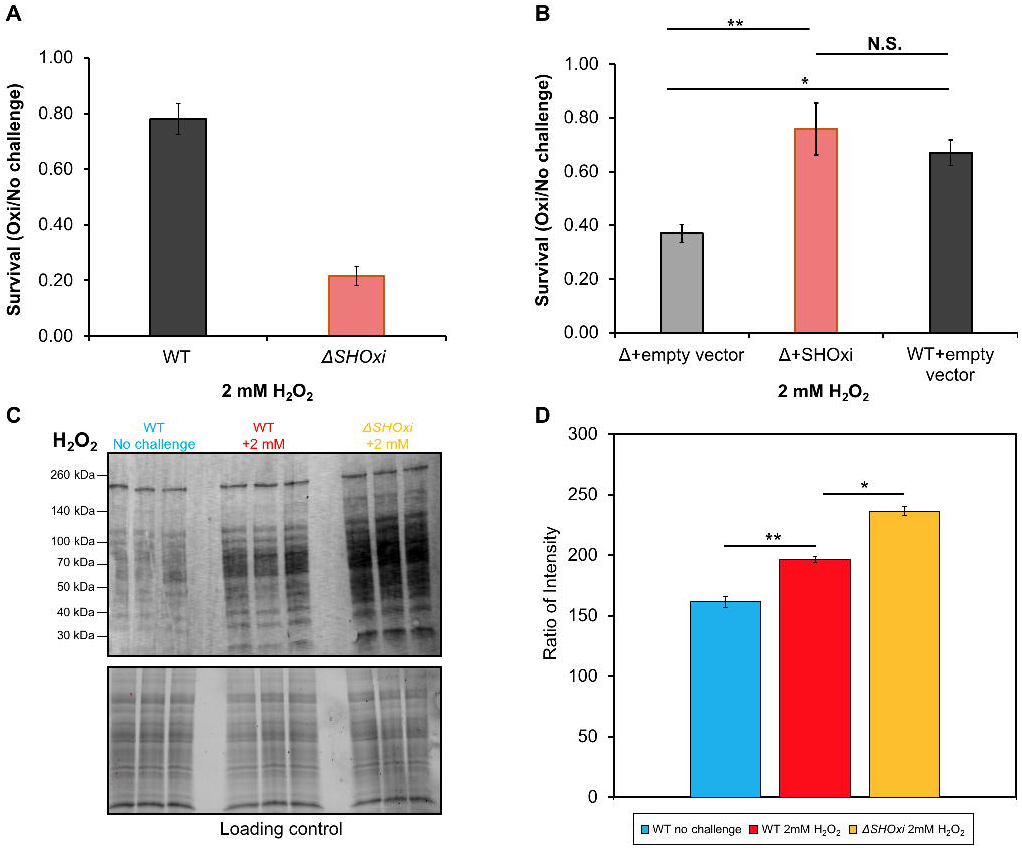
Phenotypic characterization of *ΔSHOxi*. (A) Survival of wild type and *ΔSHOxi* under oxidative stress. (B) Rescuing survival by overexpression of SHOxi in a *ΔSHOxi* mutant. The negative control was *ΔSHOxi* with an empty vector (Δ + empty vector), and the positive control was the wild type with an empty vector (WT+empty vector). SHOxi was overexpressed on the plasmid pTA1300 under a tryptophan inducible promoter in a *ΔSHOxi* background. In both (A) and (B), survival was calculated as the ratio of colony forming units (CFU) between no challenge and oxidative stress conditions (± 2 mM H_2_O_2_, 1h exposure). (C) Western blot analysis of carbonyl groups found on proteins, a proxy for oxidative damage. Samples were the wild type under no challenge conditions (0 mM H_2_O_2_), the wild type exposed to 2 mM H_2_O_2_ (80% survival), and *ΔSHOxi* exposed to 2 mM H_2_O_2_. A loading control gel is provided below the plot. (D) Quantification of (C) with corresponding legends.

**FIGURE 3:**
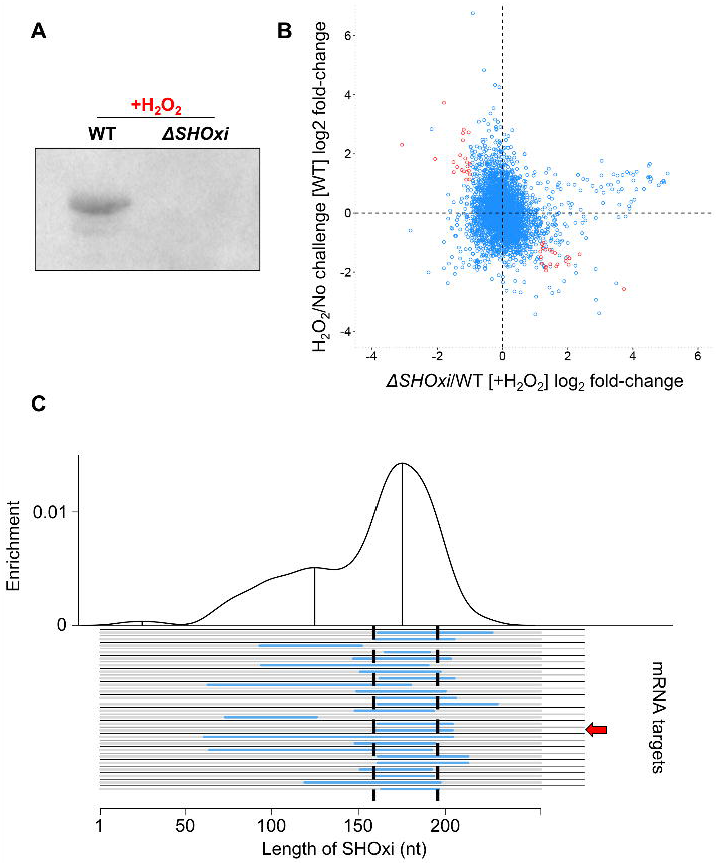
Identification of potential targets of SHOxi. ((A) Northern blot analysis confirming the presence SHOxi transcripts in WT and absence in *ΔSHOxi*. (B) Scatterplot of differentially expressed genes in the absence of SHOxi (fold-change between *ΔSHOxi* +H_2_O_2_ and WT +H_2_O_2_) compared to the wild type under oxidative stress (fold-change between +H_2_O_2_ and no challenge). (C) Predicted hybridization plot of the top 25 most probable interactions in the entire transcriptome of *H. volcanii.*

To demonstrate whether SHOxi was directly involved in cell survival during oxidative stress, we constructed an overexpression strain with the SHOxi gene under an inducible tryptophan promoter (pTA1300) (**Fig. S5A**). Using RNA-seq, we found a ~32x fold increase in SHOxi mRNA levels in both the no challenge and oxidative stress conditions, relative to WT under oxidative stress (**Fig. S5B**). WT *H. volcanii* transformed with an empty vector yielded ~ 67% survival during oxidative stress (**Fig. 2B**, +2 mM H_2_O_2_, 1h) whereas *ΔSHOxi* transformed with an empty vector yielded low survival (avg. 37% survival) under the same conditions (**Fig. 2B**). However, ectopic expression of SHOxi in a *ΔSHOxi* background resulted in rescued survival levels (avg. 76% survival) under oxidative stress conditions, comparable to WT (**Fig. 2B**), and establishing that SHOxi was directly involved in the survival of *H. volcanii* during oxidative stress.

To test whether the observed decrease in survival of the SHOxi deletion mutant under oxidative stress was related to increased oxidative damage to the cell’s macromolecules, we used Western blot analysis to measure the level of protein carbonylation. The addition of carbonyl groups to side chains of amino acid residues (lysine, arginine, proline, and threonine) is an irreversible oxidative damage that can be used as a proxy to measure levels of oxidative damage in cells (48). We found an increase in carbonyl groups in the WT under oxidative stress compared to no challenge conditions, indicating oxidative damage to proteins as a result of H_2_O_2_ exposure (**Fig. 2C-D**, WT +2mM H_2_O_2_). When measuring protein carbonylation in *ΔSHOxi* under oxidative stress, we found an additional increase of ROS-mediated damage (~1.64X) compared to WT under oxidative stress (**Fig. 2C-D**, *ΔSHOxi* +2mM H_2_O_2_). This indicated that SHOxi was likely involved in modulating the level of oxidative damage to macromolecules in the cell, which in turn, impacted the survival of *H. volcanii* during oxidative stress. (**Fig. 2D**).

### SHOxi alters the expression of putative target mRNAs

SHOxi is an intergenic sRNA and, as such, has incomplete complementarity to its mRNA targets, making it difficult to find putative targets by sequence analysis. To identify SHOxi targets, we sequenced the transcriptome of *ΔSHOxi* and WT *H. volcanii* under oxidative stress conditions, and WT *H. volcanii* under no challenge conditions. We found 215 mRNAs with significant log_2_ fold-changes (≥2) and with a false discovery rate less than 5% between *ΔSHOxi* and WT during oxidative stress (**Fig. 3B**, **Table S1**). We further restricted these putative targets to only include mRNAs with opposite fold change patterns (≥2) between WT oxidative stress and WT no challenge conditions, to increase the stringency of our analysis and the potential of selecting mRNA targets affected only by SHOxi and not by other factors (i.e. oxidative stress). Using these stringent criteria, 46 putative targets of SHOxi were identified (**Fig. 3B**, **red dots, Table S1**). A gene ontology analysis (DAVID) found that putative targets up-regulated in the absence of SHOxi were significantly (p<0.05) enriched for transcriptional regulators, while down-regulated targets were enriched for sugar metabolism.

An *in-silico* approach was also used to (i) find interacting partners based on sRNA-mRNA hybridization interactions and (ii) determine whether there was a region within SHOxi most probable for these interactions (i.e. lowest free energy change). We used IntaRNA to calculate hybridization energies between SHOxi and all the transcripts in the NCBI *H. volcanii* genome annotation. This analysis yielded a 20 nt conserved region in SHOxi that was putatively assigned as the interaction site for the 25 most reliably predicted targets (p<0.01) (**Fig. 3C**, **Table S1**). This putative interaction site corresponded to a multi stem-loop region in the modeled secondary structure of SHOxi (**Fig. 1E**, **magenta**).

By intersecting our *in-silico* and experimental approaches to identify SHOxi targets, we found one transcript mRNA that was significantly up-regulated (FDR = 5.48E-12) in *ΔSHOxi* and was predicted to have strong RNA-RNA interaction with SHOxi (**Fig. 3C**, **red arrow**). This mRNA was annotated as a bifunctional malic enzyme oxidoreductase/phosphotransacetylase (HVO_RS16435). Malic enzyme is an enzyme of the tricarboxylic acid (TCA) cycle that catalyzes the oxidative decarboxylation of malate into pyruvate using NAD^+^ or NADP^+^ as cofactors (49). SHOxi was predicted to interact with a 40 nt region, ~300 nt downstream of the TSS of malic enzyme mRNA, with a significantly strong hybridization energy (−31 kcal/mol, p = 0.00128) (**Fig 4A**). The region of interaction corresponded to a putative “seed” region (a segment of contiguous base-pairing) at 166 to 174 nt within the predicted stem loop interaction site of SHOxi (**Fig. 4A**).

**FIGURE 4:**
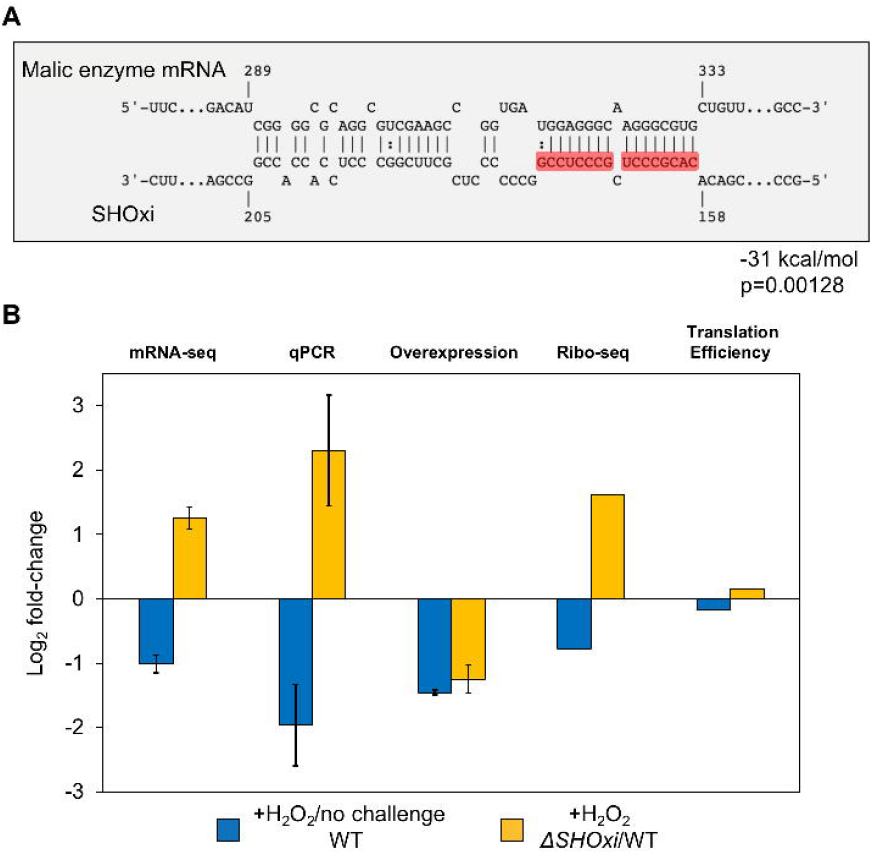
Regulation of malic enzyme mRNA by SHOxi. (A) Predicted interaction between malic enzyme mRNA (target) and SHOxi (query). Wobble base bonds indicated by two dots between hybridizing nucleotides. (B) Bar plot of log_2_ fold-change of malic enzyme mRNA from mRNA-seq, qPCR, and SHOxi overexpression, and translation efficiency of malic enzyme mRNA (Ribo-seq).

### The stem loop region of SHOxi directly interacts with malic enzyme mRNA to regulate its expression

In our RNA-seq dataset, malic enzyme mRNA was in the top 11^th^ percentile of mRNA expression levels during no challenge conditions, indicating that this transcript was one of the most highly expressed in *H. volcanii*. Under oxidative stress, malic enzyme mRNA levels decreased 2-fold when compared to no challenge condition, dropping to the 30^th^ percentile of mRNA expression levels (**Fig. 4B)**. In contrast, in *ΔSHOxi*, malic enzyme mRNA increased more than 2-fold in when compared to the WT under oxidative stress (**Fig. 4B**). This differential regulation combined with a significant *in silico* binding interaction suggested that malic enzyme mRNA might be a direct target of SHOxi. We validated our RNA-seq results by using quantitative (q)RT-PCR in WT, *ΔSHOxi*, and in our constructs overexpressing SHOxi in a WT or *ΔSHOxi* background, under oxidative stress (+2 mM H_2_O_2_, 1h) (**Fig. 4B**). We then hypothesized that while the alteration of malic enzyme transcript levels may play a role in the decreased survival of *H. volcanii*, translation of the transcript might also be affected. We recently developed ribosome profiling in *H. volcanii* (50), a global measure of translation in a cell, and carried out ribosome profiling on WT and *ΔSHOxi* under no challenge and oxidative stress conditions. We found that malic enzyme mRNA translation levels correlated with transcription levels (**Fig. 4B**) and that translation efficiency measurements were not significantly different between WT and *ΔSHOxi* under oxidative stress (**Fig. 4B**). These results indicated that SHOxi’s regulatory effect on malic enzyme mRNA was most likely upstream of translation and that it was potentially mediated post-transcriptionally by direct RNA-RNA interactions.

To establish that there was an in *vivo* binding interaction between SHOxi and malic enzyme mRNA, we used site-directed mutagenesis to construct 7 SHOxi mutant constructs with various mutations in the predicted “seed” binding region (**Fig. 5A**).The SHOxi mutant constructs were experimentally tested by ectopic overexpression in a *ΔSHOxi* background using an inducible tryptophan promoter. In the absence of tryptophan, we found little to no expression of SHOxi whereas in the presence of tryptophan there was a drastic increase in SHOxi expression level (**Fig. 5B**, +2 mM tryptophan for 1h). Malic enzyme mRNA levels were measured using qRT-PCR for each of the SHOxi mutants and fold changes were calculated relative to the overexpression of the non-mutated SHOxi in a *ΔSHOxi* background. We found that the *ΔSHOxi* background strain with an empty vector (**Fig. 5C**, Empty vector) had higher malic enzyme mRNA levels (~2x) than the *ΔSHOxi* strain overexpressing a non-mutated SHOxi (**Fig. 5C**, WT OE). Most point or dinucleotide mutations in the “seed” binding region of SHOxi did not alter expression of malic enzyme mRNA (**Fig. 5C**, Mut1, Mut2, Mut 5, and Mut 6), indicating a non-disruptive effect on the binding between the two transcripts. In contrast, a point mutation at position 163 in SHOxi, from a guanine to a cytosine (**Fig. 5C**, Mut3) or an adenosine (**Fig. 5C**, Mut4), increased malic enzyme mRNA expression by 2 and 2.2 fold, respectively. Lastly, a combination of all 5 nucleotide mutations in the “seed” binding region of SHOxi (**Fig. 5C**, Mut7) resulted in an increase of malic enzyme mRNA levels comparable to that of the empty vector. These results confirmed our prediction for an *in vivo* binding interaction between malic enzyme mRNA and the stem loop region of SHOxi and showed that mutation at position 163 in SHOxi had a strong disruptive effect on this interaction (**Fig. 5C**).

**FIGURE 5:**
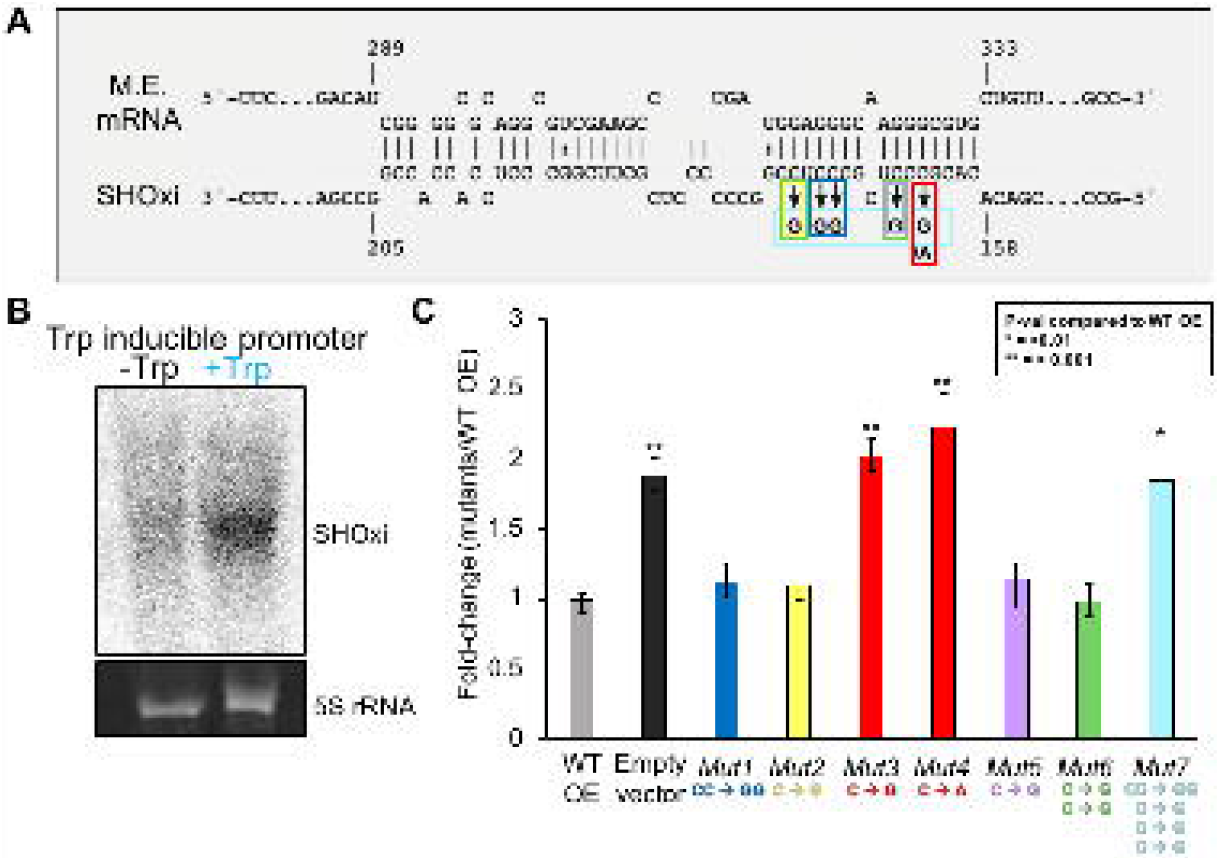
*In vivo* validation of direct interaction between SHOxi-malic enzyme mRNA. (A) Schematic of mutations in the “seed” binding region of SHOxi with malic enzyme mRNA. (B) Northern blot of SHOxi transcript levels expressed ectopically under a tryptophan promoter. -Trp is without tryptophan, and +Trp is with 2mM tryptophan for 1h. 5S rRNA is a loading control. (C) qRT-PCR of malic enzyme mRNA levels in various SHOxi mutants (from A) calculated as the fold change relative to overexpression of the WT SHOxi.

### Malic enzyme mRNA stability is post-transcriptionally regulated by SHOxi

To further investigate how SHOxi directly affected malic enzyme transcript levels, we measured malic enzyme mRNA stability *in vivo* following the addition of H_2_O_2_ to induce SHOxi expression (**Fig. 6A**). After 30 min, Actinomycin D (actD) was added to WT and *ΔSHOxi* cultures to inhibit transcription, total RNA was extracted at time intervals of 0, 15, 30, and 60 min after actD addition, and malic enzyme mRNA levels were measured using qPCR (**Fig. 6A**). A house keeping gene, the surface glycoprotein, with no altered expression level in WT and *ΔSHOxi*, was used as a control. We found that malic enzyme mRNA transcript levels were higher over the time course in *ΔSHOxi* compared to WT under oxidative stress, indicating that the mRNA was more stable in the absence of SHOxi (**Fig. 6B**). In contrast, the surface glycoprotein mRNA did not show any significant difference in transcript levels between *ΔSHOxi* and WT under oxidative stress (**Fig. 6C**).

**FIGURE 6:**
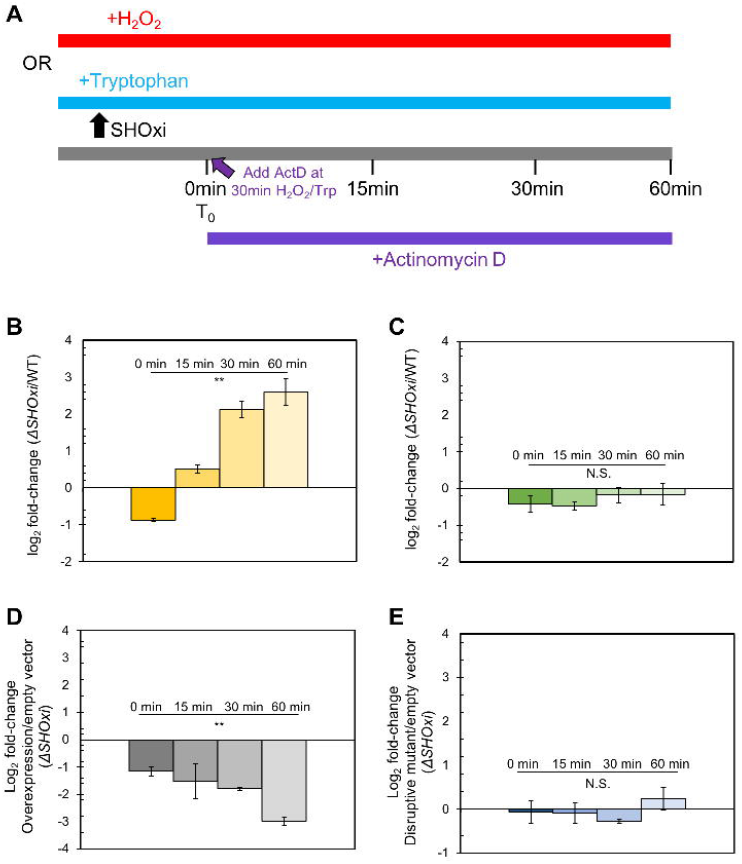
Destabilization of malic enzyme mRNA. (A) Schematic of the experimental approach to measure the half-life of malic enzyme mRNA in response to SHOxi. 30 min after SHOxi induction, with H_2_O_2_ or via an inducible tryptophan promoter, transcription was shut off by the addition of 100 μg/ml actinomycin D and the level of malic enzyme mRNA was analyzed by qRT-PCR at several time points. (B) qRT-PCR of malic enzyme mRNA levels over time after addition of actinomycin D. Log_2_ fold changes were calculated between the WT and *ΔSHOxi* with 2 mM H_2_O_2_. (C) qRT-PCR of the house keeping gene surface protein mRNA levels in the same conditions as in (B). (D) qRT-PCR of malic enzyme mRNA levels after addition of actinomycin D and the induction of SHOxi overexpression with 2 mM tryptophan, under no challenge conditions. Log_2_ fold changes were calculated between the SHOxi overexpression construct and the empty vector in a *ΔSHOxi* background. (E) qRT-PCR of malic enzyme mRNA levels in the same conditions as in (D) but with the overexpression of the disruptive SHOxi mutant Mut 7 from **Fig. 4–5C**.

Next we repeated the experiment this time driving the ectopic overexpression of SHOxi with tryptophan, in a *ΔSHOxi* background (**Fig. 6 D-E**). We found that malic enzyme mRNA transcript levels were lower over time with SHOxi overexpression compared to the empty vector control (**Fig. 6 D**). In contrast, overexpressing a disruptive SHOxi mutant (Mut 7 from **Fig. 5C)** did not result in a significant difference in malic enzyme mRNA transcript levels compared to the empty vector (**Fig. 6 E**). These findings clearly showed that malic enzyme mRNA was destabilized by SHOxi during oxidative stress.

### Malic enzyme modulates the ratio of NAD+/NADH during oxidative stress

As noted before, malic enzyme is bifunctional in its usage of dinucleotides in the oxidation of malate to pyruvate and CO_2_ in central metabolism (49, 51, 52). Certain variants of malic enzyme are NAD^+^-dependent, converting NAD^+^ to NADH, while other variants are NADP^+^-dependent, converting NADP^+^ to NADPH (49, 51). In Eukaryotes malic enzymes are highly conserved and use NADP^+^ while both enzyme variants are found in prokaryotes (49). For example, *E. coli* encodes two variants of malic enzyme in its genome; SfcA is C-terminally truncated and NAD+-dependent while MaeB is longer and NADP^+^-dependent (49). *H. volcanii* has two variants of malic enzyme (**Fig. S6**), one of which was regulated by SHOxi (HVO_RS16435) while the other (HVO_RS15075) was not nor did its expression level change during oxidative stress. The two variants shared high sequence identity at the protein level (**Fig. S6A**) and their co-factor specificity was derived from genomic annotations. Using I-TASSER, we generated high confidence protein models for both *H. volcanii* malic enzymes (**Fig. S6B-D**). Ligand binding predictions with these protein models showed that both malic enzymes had a high confidence malate-binding domain (**Fig. S6B**). We also found that HVO_RS16435, the malic enzyme variant regulated by SHOxi, only had a binding domain for NAD^+^ while HVO_RS15075, not regulated by SHOxi, had both a high confidence NADP^+^ and low confidence NAD^+^ binding domain (**Fig. S6B**).

Biochemical analyses are inherently limited in *H. volcanii* due to the high intracellular salt dependency of the proteome (27). Instead, to elucidate the cofactor-binding capacity of HVO_RS16435, the SHOxi-regulated malic enzyme, we measured all nicotinamide adenine dinucleotides in the cell (NAD+, NADH, NADP+, NADPH) using a luciferase-based assay (Promega) for both WT and *ΔSHOxi* during no challenge and oxidative stress conditions. We then calculated ratios between NAD^+^:NADH and NADP^+^:NADPH levels to normalize the results between different conditions and strains. We found that the NAD^+^:NADH ratio in the WT increased under oxidative stress (when SHOxi is up-regulated) relative to the WT in no challenge conditions (**Fig. 7**). In contrast, the NAD^+^:NADH ratio decreased in *ΔSHOxi* compared to WT under oxidative stress. The ratio of NADP^+^:NADPH was not altered for any of the experimental conditions (**Fig. 7**), indicating that the function of the SHOxi-regulated malic enzyme (HVO_RS16435) is likely NAD^+^-dependent.

**FIGURE 7:**
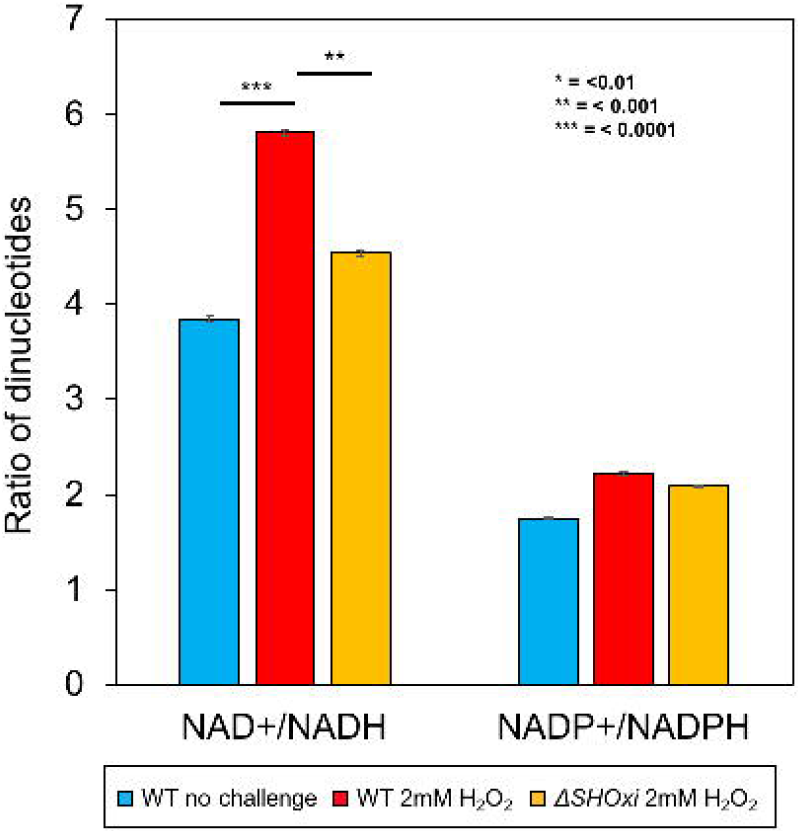
Measurements of cellular dinucleotides using a luciferase-based assay. Ratios of NAD+/NADH and NADP+/NADPH were calculated to determine the relative abundance.

## DISCUSSION

By elucidating the mechanism and function of the most up-regulated intergenic sRNA, SHOxi, during H_2_O_2_-induced oxidative stress in *H. volcanii,* we showed that SHOxi was a major regulator of the oxidative stress response. Under oxidative stress, SHOxi was enriched 21-fold and its deletion (*ΔSHOxi*) resulted in a drastic increase in oxidative damage to the cell’s macromolecules and a severe decrease in *H. volcanii* survival, underlying its key role in the stress response.

In Bacteria, several sRNAs have been implicated in the regulation of the oxidative stress response (37, 38, 41, 53). The bacterial sRNA OxyS is the most studied of those and its regulatory mechanism has recently been fully characterized (38). OxyS is implicated in protecting *Escherichia coli* cells from DNA damage by decreasing the translation of the essential transcription termination factor NusG. This leads to an increase of the virulence factor kilR, which, in turn, interferes with the cell division protein FtsZ and ultimately inhibits cell division. The arrest in cell growth provides more time for DNA damage repair, hence better survival under oxidative stress (38).

Using mRNA-seq, we identified several putative targets that were differential expressed in presence (WT) and absence (*ΔSHOxi*) of SHOxi under oxidative stress and no challenge conditions. These putative targets included several transcription factors with yet unknown functions, suggesting that SHOxi may be a master regulator with large downstream effects in the gene regulatory network. Interestingly, these transcription factor mRNAs had increased transcript levels in the presence of SHOxi, indicating that they might be stabilized by the sRNA. Several other putative targets were down-regulated in the presence of SHOxi, including a sugar ABC transporter operon and malic enzyme. This is not surprising since dual functioning sRNAs have been reported in other Archaea, such as sRNA_154_ involved in nitrogen metabolism in *M. mazei* (13, 25).

Here we report on one specific target of SHOxi, malic enzyme, and present strong experimental evidence that SHOxi mediates the degradation of malic enzyme mRNA under oxidative stress. Malic enzyme mRNA was highly expressed in *H. volcanii* under no challenge condition and was significantly down-regulated in the presence of SHOxi. We demonstrated that SHOxi binds to malic enzyme mRNA *in vivo* with strong RNA-RNA interaction and extensive base pairing of the stem-loop region in SHOxi’s secondary structure. By measuring steady-state RNA levels, we also showed that the stability of malic enzyme mRNA decreased over time only when SHOxi was present in the cell. These findings strongly support a mechanism by which SHOxi regulate malic enzyme during oxidative stress by destabilizing its mRNA through direct RNA-RNA binding interactions.

Destabilization of mRNAs through RNA-RNA interactions by sRNAs and the recruitment of a RNase has been well documented in bacteria (54, 55). We speculate here that the destabilization of malic enzyme mRNA by SHOxi is the result of the activity of a currently unknown RNase. While RNases and RNA degradation pathways are not well resolved in Archaea, recent studies have brought insights onto potential biochemical mechanisms for novel RNases (56– 60). Both endonucleases and exonucleases have been reported in the archaea but because of the internal interacting site of the malic enzyme mRNA with SHOxi, we speculate that an endonuclease is the most likely RNase candidate. Intriguingly, a RidA endonuclease was found to degrade the mRNA of a potassium transporter in response to shifts in extracellular potassium concentrations in *H. salinarum* (58). The corresponding RidA homolog in *H. volcanii* is indeed the most up-regulated gene during oxidative stress, which may indicate a key role in RNA processing during oxidative stress (9).

Malic enzymes decarboxylate L-malate to pyruvate using either NAD^+^/NADH or NADP^+^/NADPH as co-factors. These enzymes regulate metabolic flux in central carbon metabolism by linking glycolysis and gluconeogenesis with the TCA cycle (49, 51, 52). *H. volcanii* encodes two variants of malic enzymes, similarly to *E. coli*, with one variant regulated by SHOxi (HVO_RS16435) while the other is not (HVO_RS15075). Using protein modeling, we demonstrated that both malic enzymes had a high confidence malate-binding domain but that the HVO_RS16435 variant had a putative NAD^+^-binding domain while the other variant had a putative NADP^+^-binding domain. Measuring nicotinamide adenine dinucleotides ratios in *H. volcanii,* in presence or absence of SHOxi and under oxidative stress or no challenge conditions, confirmed the NAD^+^/NADH-specificity of the HVO_RS16435 malic enzyme variant. We therefore argue that the increase of the NAD^+^/NADH ratio we observed under oxidative stress was the functional consequence of SHOxi-mediated post-transcriptional regulation of the HVO_RS16435 malic enzyme mRNA. Whether this was a direct effect by malic enzyme on the NAD^+^/NADH ratio or via the downregulation of the TCA cycle remains to the demonstrated. Of interest, previous studies with haloarchaea, and this work (**Fig. S7**), have shown that central metabolism was downregulated under oxidative stress, including all the enzymes of the TCA cycle, potentially resulting in a decrease production of NADH, a pro-oxidant, in a cell trying to maintain a reducing environment and minimize the production of ROS (9, 61, 62). Indeed, NADH, generated primarily in the TCA cycle, is oxidized at the electron transport chain (ETC). Electrons from this oxidation are shuttled along the ETC to ultimately reduce oxygen to water in a process coupled with the generation of a proton gradient and ATP synthesis (63). However, the ETC is prone to leakage, generating superoxide, as an inevitable by-product of the activity of Complexes I and III, and the production of other ROS in the cell (64).

The regulation of reduced nicotinamide nucleotides NADH and NADPH under oxidative stress has been reported in other microorganisms (65, 66). In *Pseudomonas fluorescence* NADPH-generating enzymes were highly upregulated during menadione-induced oxidative stress while NADH-generating enzymes were down regulated (66). Moreover, NAD^+^ kinase and NADP^+^ phosphatase, enzymes that regulate the levels of NAD^+^ and NADP^+^, showed altered activity during oxidative stress, promoting a reducing intracellular milieu (less NAD^+^, more NADP^+^). While we propose here that *H. volcanii* is actively decreasing the NAD^+^/NADH ratio via the SHOxi-mediated regulation of malic enzyme under oxidative stress, it is important to consider that non-SHOxi mediated metabolic shifts (such as in *P. fluorescences*) might also be part of the cell’s response to H_2_O_2_ treatment.

We propose a model for SHOxi-mediated regulation of malic enzyme mRNA (**Fig. 8**). In no challenge condition, and the absence of SHOxi, malic enzyme mRNA is highly expressed, NADH is generated via the TCA cycle for ATP production via oxidative phosphorylation, and redox homeostasis is maintained by a balanced ratio of NADH/NADPH. (**Fig. 8A**). Under oxidative stress, SHOxi is drastically up-regulated, resulting in the destabilization of the malic enzyme mRNA via base pair interactions with SHOxi and a yet unknown RNase, and the NAD+/NADH ratio is increased compared to no challenge condition (**Fig. 8B**). Whether this interaction is aided by a protein chaperone, such as Hfq in Bacteria, is not known. The reduced oxidative damage to the cell’s macromolecules from the activity of SHOxi results in a moderate decrease in survival (**Fig. 8B**). In a SHOxi knockout mutant (*ΔSHOxi*), where there is no production of SHOxi even during oxidative stress, the malic enzyme mRNA remain highly expressed and the NAD+/NADH ratio is similar to that of no challenge condition (and lower than in the presence of SHOxi) (**Fig. 8C**). As a consequence, an increase in oxidative damage to the cell is observed, and survival is severely reduced (**Fig. 8C**). The increased ratio of NAD^+^/NADH in presence of SHOxi indicates that the survival of *H. volcanii* to oxidative stress may, in part, be linked to the role malic enzyme plays in the TCA cycle and the interplay of dinucleotides generation for redox homeostasis (**Fig. 8B-C**). Alternatively, an increase in cellular NAD^+^, as the result of SHOxi activation under oxidative stress, could provide an additional template for the enzymatic conversion of NAD^+^ to NADP^+^ by an NAD kinase, generating NADPH, a strong antioxidant (67, 68). However, we did not find evidence for a change in the ratio of NADP^+^/NADPH under oxidative stress conditions in *H. volcanii*.

**FIGURE 8:**
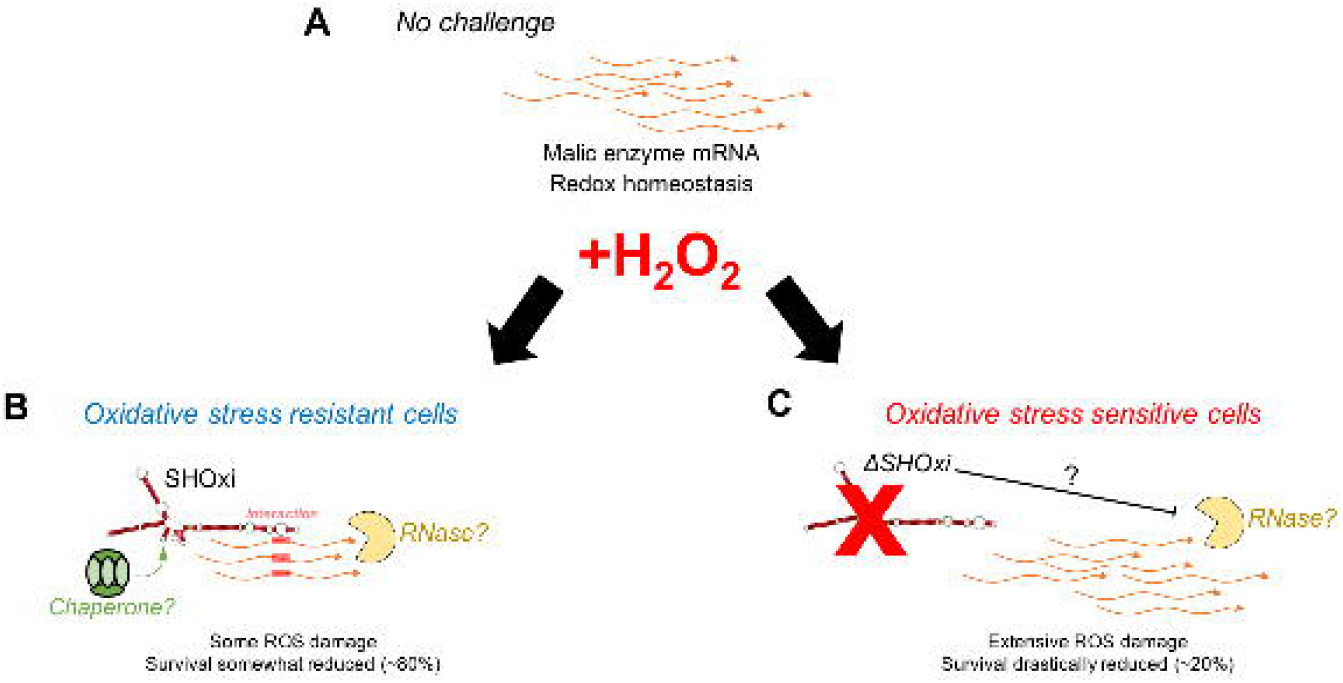
A model for the posttranscriptional regulation of malic enzyme mRNA by SHOxi. (A) Wild type no challenge conditions. (B) Wild type under oxidative stress conditions (+2 mM H_2_O_2_, 80% survival). (C) A knockout of SHOxi (*ΔSHOxi*) under the same conditions as (B).

It is clear that SHOxi-mediated destabilization of malic enzyme mRNA is just one component of the oxidative stress response of *H. volcanii.* Previous work in *H. volcanii* and *H. salinarum* has shown that oxidative stress impacted a wide array of cellular processes, engaging at least 50% of all genes (9, 35, 61). These changes were characterized by the up-regulation of DNA repair enzymes (*Rpa*A/B genes), ROS scavenging enzymes (e.g. catalase), and iron sulfur assembly proteins, high protein turnover, and the down-regulation of metabolism (9, 31, 33, 69, 70). In *H. salinarum*, but not *H. volcanii,* the transcription factor RosR was found to regulate hundreds of genes during oxidative stress (31).

In addition to malic enzyme, we found other putative targets of SHOxi in our RNA-seq screen, including transcription factors that were positively regulated in the presence of SHOxi. This suggest that SHOxi may be integrated in a complex gene regulatory network as part of the oxidative stress response in *H. volcanii*, where a transcription factor regulates SHOxi expression and SHOxi then further regulates other transcription factors for downstream gene regulation. Indeed, in both eukarya and bacteria, sRNAs have been shown to be integrated in gene regulatory networks along with transcription factors (71). The potential sRNA-mediated regulation of redox homeostasis via the modulation of NAD^+^/NADH ratio could be part of the global cellular response to oxidative damage that includes the upregulation of specific enzymatic detoxification systems (superoxide dismutases, catalases, peroxidases) and antioxidants (glutathione) (72) and the down-regulation of metabolism, as previously been reported in haloarchaea (9, 31, 33, 69, 70). Future work will include answering questions regarding the functional role of other (non-malic enzyme) SHOxi targets, which RNase might be involved in the destabilization mechanism, and whether RNA binding proteins help facilitate the interactions between SHOxi and its mRNA targets.

## CONTRIBUTIONS

DRG: Conceptualization, Investigation, Methodology, Project administration, Writing – original draft, Writing – review & editing

RR: Investigation – Sample acquisition, genetics.

KW: Investigation – Sample acquisition, genetics.

SD: Investigation – Sample acquisition, genetics, qPCR.

JDR: Conceptualization, Funding acquisition, Project administration, Supervision, Validation, Writing – review & editing

## ACKNOWLEDGEMENTS

This work was supported by NASA grant 18-EXO18-0091 and Air Force Office of Scientific Research grant FA9550-14-1-0118. We thank Kate Huffer for help in generating SHOxi deletion mutants. We thank Katie Farney for extensive manuscript editing. We thank David Mohr for sequencing technical supports, and Drs. John Kim, Gisela Storz, Sarah Woodson, and Emine Ertekin for experimental advice and helpful discussions.

## SUPPLEMENTAL FIGURE LEGENDS

**FIGURE S1:** ORF potential plots for the six reading frames of SHOxi, the protein coding mRNA RidA, and the non-coding RNA signal recognition particle.

**FIGURE S2:** Reliability plot of the secondary structure of SHOxi. Reliability scores are calculated based off of either structure conservation or sequence conservation across the length of the sRNA in the context of its secondary structure.

**FIGURE S3:** (A) PCR validation of SHOxi deletion in the genome of *H. volcanii.* WT denotes wild type genomic DNA and Δ denotes *ΔSHOxi* genomic DNA. Primers targeted 500 bp upstream and downstream of SHOxi and the expected product size was 1.5 kb. (B) RT-PCR validation of SHOxi transcript depletion levels. +, wild type genomic DNA (positive control); WT, wild type cDNA; Δ, *ΔSHOxi* cDNA. Primers were 100 bp internal to SHOxi and the expected product size was 100 bp. Multiple constructs were tested for (A) and (B).

**FIGURE S4:** Growth curves of WT and *ΔSHOxi* under (A) no challenge and (B) oxidative stress conditions. Oxidative stress induction (500 uM) is indicated by the red arrow. Unpaired t-test p-values are given for the points along the growth curve that show a difference.

**FIGURE S5:** (A) Plasmid map for expression vector pTA1300 constructed for the overexpression of SHOxi under an inducible tryptophan (trp) promoter. (B) Log_2_-fold changes of SHOxi overexpression from pTA1300 relative to wild type SHOxi endogenous expression under oxidative stress measured by RNA-seq. OE ctrl is the overexpression of wild type SHOxi under no challenge conditions and OE oxi is the overexpression of wild type SHOxi under oxidative stress conditions.

**FIGURE S6:** Protein modeling to elucidate the functions of malic enzyme mRNA variants. HVO_RS16435 is variant 1 and HV_RS15075 is variant 2. (A) Multiple sequence alignment of the protein sequences for the two malic enzyme variants. (B) Protein models using I-TASSER for the two malic enzyme variants. The first columns of panels show the most confident protein model. The other panel columns are ligand-binding predictions for the protein models. Confidence scores are given for each model (TM-score and C-score, respectively). (C) A ranked list of most probable Protein Data Bank (PDB) hits for variant 1 (HVO_RS16435) and for (D) variant 2 (HV_RS15075). Rank indicates probability, PDB hit indicates the identifier on PDB, TM-score indicates the structural alignment confidence interval, RMSD indicates the distances in angstroms between the C-alpha atoms between residues that are structurally aligned by TM-align, IDEN indicates the fraction sequence identity in the structurally aligned region, and COV indicates the coverage of the alignment by TM-align and is equal to the number of structurally aligned residues divided by length of the query protein.

**FIGURE S7:** Heatmaps of fold-changes for genes involved in NAD, malate, pyruvate, and TCA metabolism for which malic enzyme (ME) mRNA plays a role. Fold changes were calculated between wild type and *ΔSHOxi* under oxidative stress.

