## supplementary figures for "Post-transcriptional regulation of redox homeostasis by the small RNA SHOxi in haloarchaea"

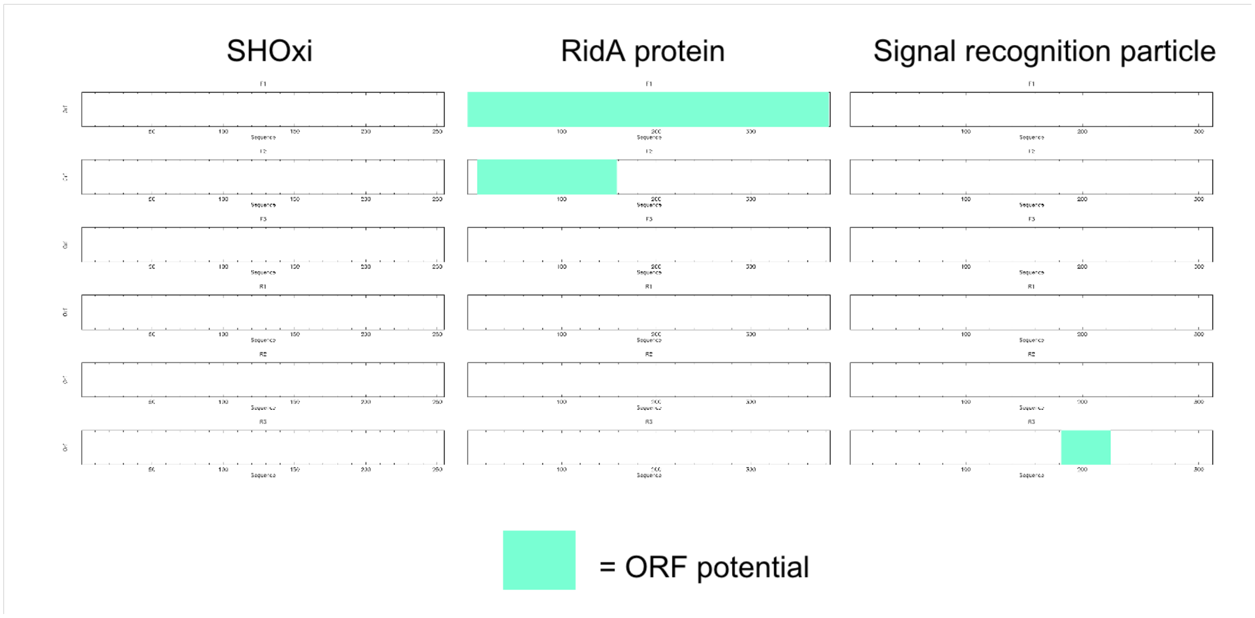


Figure S1

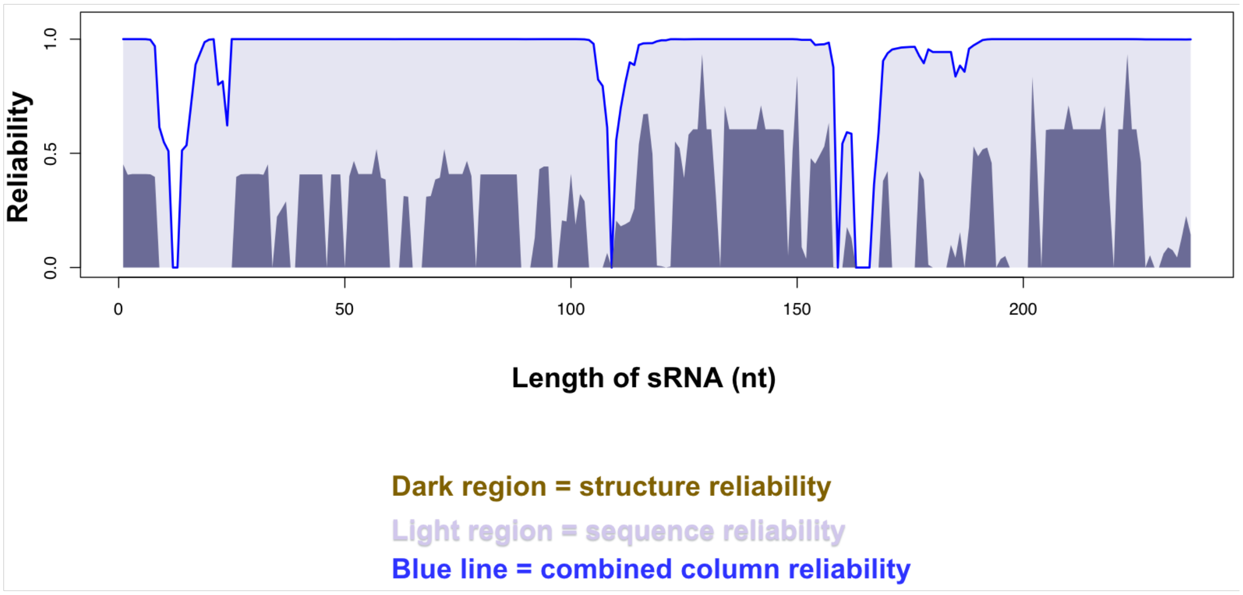

Reliability
0.0
0.5
1.0
0
50
100
150
200
Length of sRNA (nt)
Dark region = structure reliability
Light region = sequence reliability
Blue line = combined column reliability


Figure S2

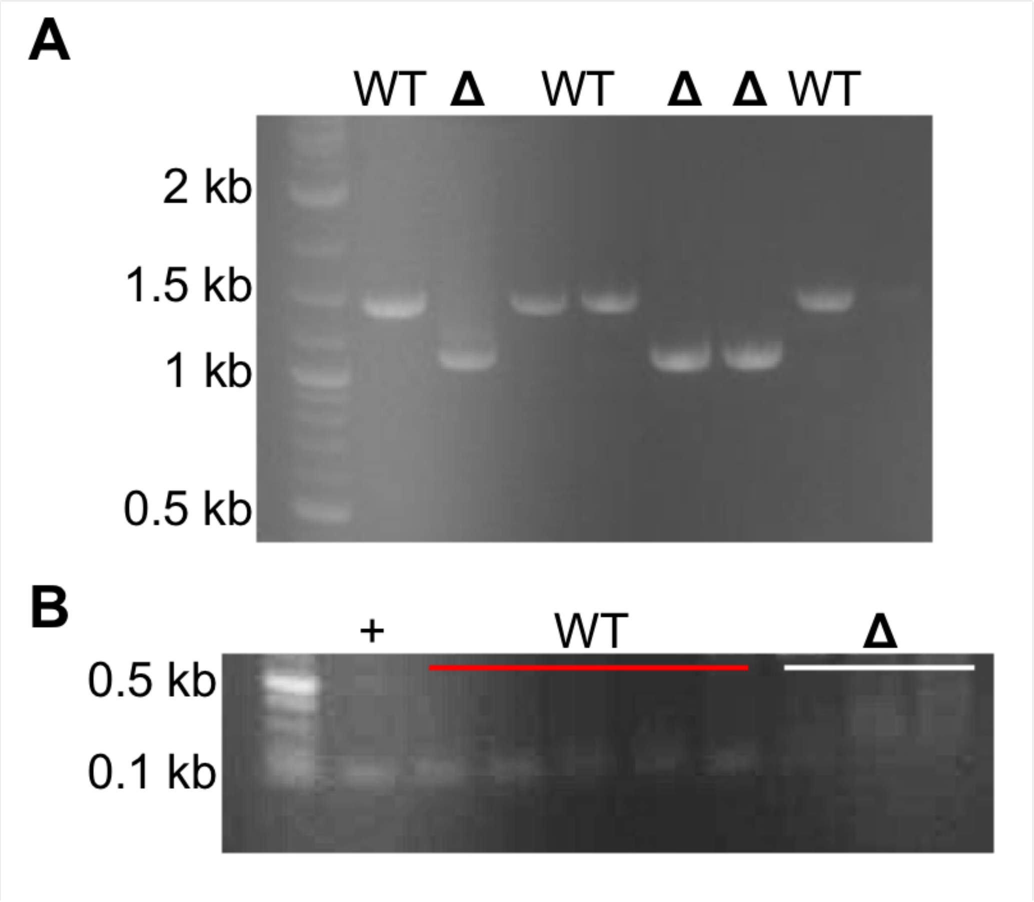


Figure S3

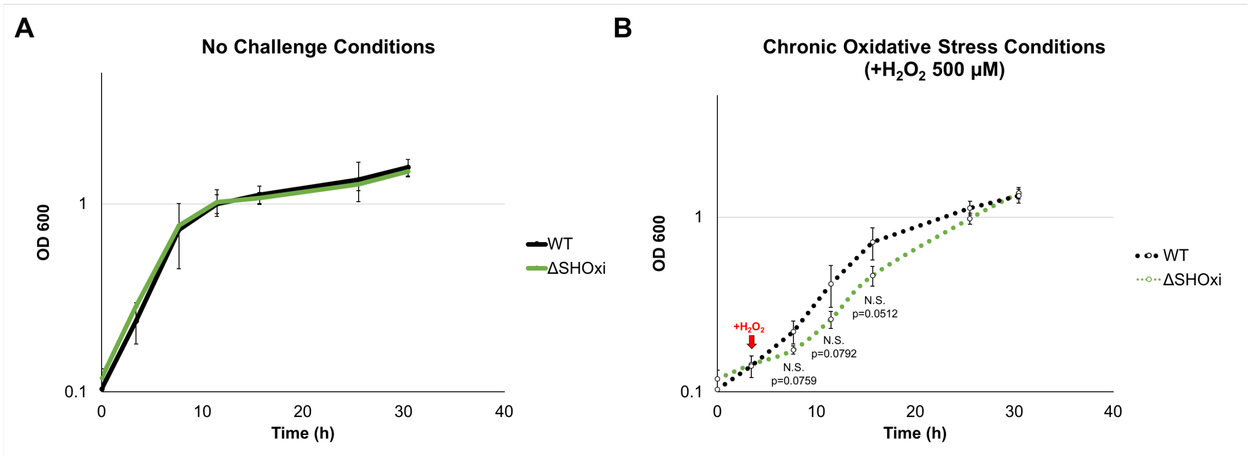
A
No Challenge Conditions
OD 600
1
0.1
0
10
20
30
40
Time (h)
WT
ΔSHOxi
B
Chronic Oxidative Stress Conditions
(+H2O2 500 μM)
OD 600
+H2O2
N.S.
p=0.0759
N.S.
p=0.0792
N.S.
p=0.0512
WT
ΔSHOxi

Figure S4

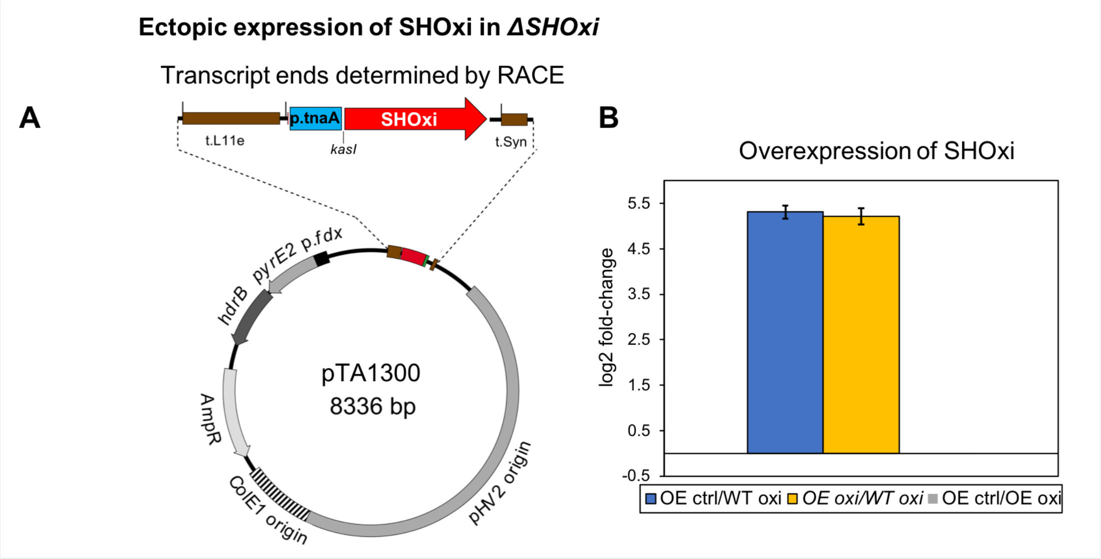
Ectopic expression of SHOxi in ΔSHOxi
Transcript ends determined by RACE
A
p.tnaA
SHOxi
t.L11e
kasI
t.Syn
pyrE2
p.fdx
hdrB
pTA1300
8336 bp
AmpR
ColE1 origin
pHV2 origin
B
Overexpression of SHOxi
log2 fold-change
5.5
4.5
3.5
2.5
1.5
0.5
-0.5
OE ctrl/WT oxi
OE oxi/WT oxi
OE ctrl/OE oxi

Figure S5

A

Two malic enzyme genes in *Hv*; **Variant1** regulated and Variant2 is not

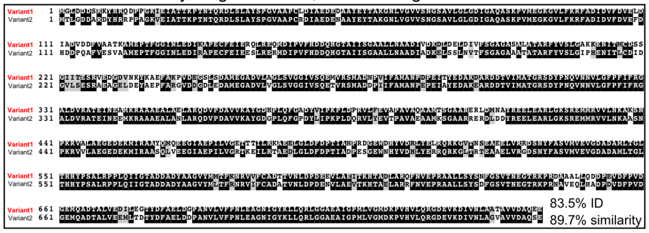


B

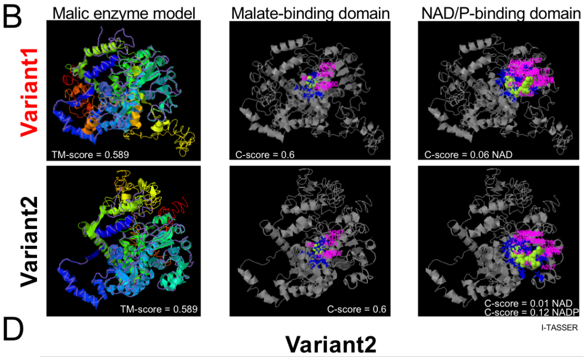


C

**Variant1**

| Rank | PDB Hit | TM-score | RMSD[a] | IDEN[a] | Cov |
|---|---|---|---|---|---|
| 1 | 3nv9A | 0.589 | 0.47 | 0.372 | 0.590 |
| 2 | 2dvmA | 0.533 | 2.52 | 0.410 | 0.565 |
| 3 | 5ceeA | 0.490 | 2.18 | 0.416 | 0.513 |
| 4 | 2haeA | 0.473 | 2.11 | 0.447 | 0.494 |
| 5 | 2a9fA | 0.466 | 2.99 | 0.421 | 0.505 |
| 6 | 1gz3A | 0.465 | 3.12 | 0.218 | 0.502 |
| 7 | 1gq2A | 0.464 | 3.20 | 0.214 | 0.503 |
| 8 | 1o0sA | 0.462 | 3.71 | 0.186 | 0.514 |
| 9 | 2aw5B | 0.456 | 3.10 | 0.205 | 0.495 |
| 10 | 5xviA | 0.349 | 5.83 | 0.065 | 0.453 |

D

**Variant2**

| Rank | PDB Hit | TM-score | RMSD[a] | IDEN[a] | Cov |
|---|---|---|---|---|---|
| 1 | 3nv9A | 0.587 | 0.57 | 0.351 | 0.589 |
| 2 | 2dvmA | 0.535 | 2.37 | 0.412 | 0.565 |
| 3 | 5ceeA | 0.491 | 2.13 | 0.435 | 0.514 |
| 4 | 2haeA | 0.473 | 2.09 | 0.442 | 0.494 |
| 5 | 2a9fA | 0.466 | 2.82 | 0.403 | 0.502 |
| 6 | 1gz3A | 0.462 | 2.94 | 0.222 | 0.495 |
| 7 | 1gq2A | 0.461 | 3.22 | 0.218 | 0.499 |
| 8 | 1o0sA | 0.456 | 3.66 | 0.190 | 0.505 |
| 9 | 2aw5B | 0.452 | 3.32 | 0.203 | 0.493 |
| 10 | 4f91B | 0.355 | 8.23 | 0.060 | 0.555 |

Figure S6

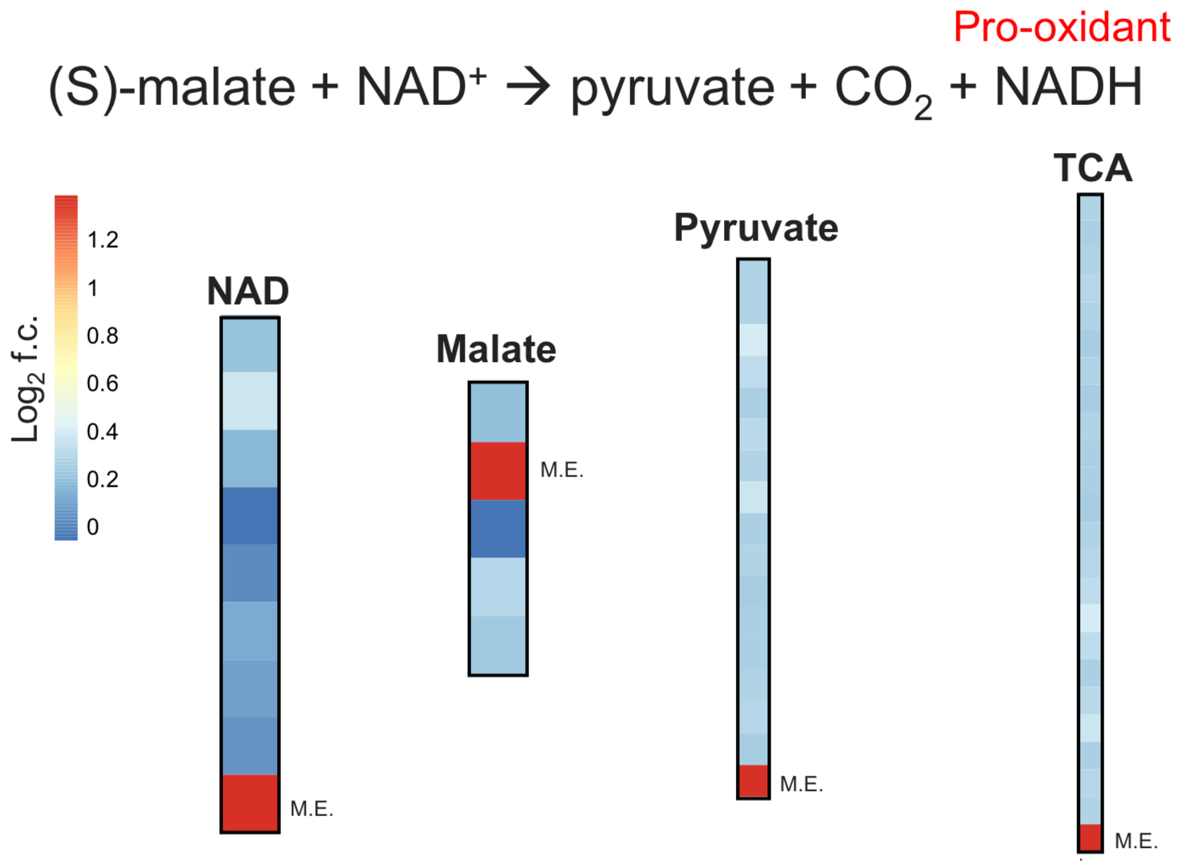
Pro-oxidant
(S)-malate + NAD+ → pyruvate + CO2 + NADH
Log2 f.c.
1.2
1
0.8
0.6
0.4
0.2
0
NAD
Malate
Pyruvate
TCA
M.E.
M.E.
M.E.
M.E.

Figure S7
